# A large language model for predicting pancreatic ductal adenocarcinoma patients from blood-derived exosomal transcriptomics data

**DOI:** 10.1101/2025.03.06.641795

**Authors:** Shubham Choudhury, Naman Kumar Mehta, Gajendra P. S. Raghava

**Affiliations:** Department of Computational Biology, Indraprastha Institute of Information Technology, New Delhi, India

**Keywords:** Transcriptomics profile, Large Language Model, Pancreatic ductal adenocarcinoma, Protein language models

## Abstract

Traditional machine learning approaches for text or sequence classification rely on converting textual data into numerical representations. In this study, we investigate a reverse strategy in which numerical features are transformed into sequence representations and classified using large language models (LLMs). We applied this methodology to predict pancreatic ductal adenocarcinoma (PDAC) using the expression profiles of 50 genes from 284 PDAC and 217 non-PDAC patients. Gene expression values were converted into sequence data, with each gene represented as a residue in a 50-residue protein sequence. Major LLMs like PeptideBERT, ProtBERT, and ESM2 were fine-tuned on a protein training dataset and evaluated on an independent dataset. The best-performing model, ProtBERT, achieved an AUC of 0.962 on an independent dataset. Additionally, an alignment-based approach employing BLAST and MERCI motifs was explored, and an ensemble model combining the LLM-based and alignment-based methods was developed. Our LLM-based model outperformed traditional machine learning models. To the best of our knowledge, this is the first study demonstrating the application of LLMs for mining transcriptomic profiles of cancer patients.

**HIGHLIGHTS:**

- Identification of over and under-expressed genes in PDAC patients
- Convert numeric gene expression data to peptide sequence
- LLM based models for predicting PDAC patients using peptide sequences
- Mining of transcriptomics data using ProtBert and ESM2
- Gene expression profile for diagnostic of PDAC patients

## INTRODUCTION

Pancreatic ductal adenocarcinoma (PDAC), which accounts for over 90% of all pancreatic tumors, is characterized by a robust inflammatory desmoplastic reaction^1^. It is the fourth leading cause of cancer-related deaths globally, with a five-year survival rate of less than 8%^2^. This dismal prognosis arises largely because PDAC is often diagnosed at advanced stages, leaving limited treatment options. Moreover, the incidence of PDAC is projected to more than double within the next decade due to factors such as population ageing, obesity, and type 2 diabetes, highlighting an urgent need for innovative and effective diagnostic strategies^3–5^. Current diagnostic approaches for PDAC rely on a combination of surgery and imaging techniques, such as computed tomography (CT), magnetic resonance imaging (MRI), and endoscopic ultrasound (EUS). However, these methods face substantial limitations, including difficulties in detecting small or early-stage tumors, dependence on operator expertise, and high costs, making them less accessible and effective. Furthermore, with only 15-20% of patients eligible for surgery and most presenting with distant metastasis, the window for meaningful intervention is narrow^6,7^.

Recognizing these challenges, researchers have sought innovative solutions, with transcriptomic markers in blood emerging as a particularly promising avenue. Despite significant progress, existing diagnostic strategies leveraging transcriptomic biomarkers—such as CA19-9 and extracellular vesicle protein markers like GPC1—often rely on traditional machine learning models like Random Forest (RF) and Lasso regression. While these methods have shown promise, they are limited in their ability to capture complex, high-dimensional patterns in gene expression data^8–11^. The emergence of the role of exosomal proteins has also been highlighted by computational methods that can accurately predict exosomal proteins^12,13^. Recent studies have demonstrated that even simple machine learning models, such as Logistic Regression and Support Vector Machines (SVM), can achieve high diagnostic accuracy when trained on curated biomarker panels^11,14–16^. In 2019, Wang et al. developed a diagnostic biomarker based on the expression of eight genes. A SVM model was trained on gene expression values calculated in transcripts per million (TPM). The eight gene biomarker signature developed in this study achieved an AUC of 0.939 on the external validation dataset, highlighting the potential of non-invasive blood-based biomarkers compared to conventional techniques.

In this study, we introduce a novel approach that employs large language models (LLMs) to analyze transcriptomic data for PDAC prediction. While LLMs have revolutionized natural language processing (NLP) by capturing intricate patterns in textual data, their application to numerical datasets has been largely unexplored. In this study, we adapted LLMs to process numerical transcriptomic features by converting them into text-based representations, enabling the models to leverage their robust pattern-recognition capabilities. We have utilized the dataset used by Wang et al. for training and validation. Our results demonstrate that the performance of LLM-based models is of a similar order to established methods, while providing a unique perspective on data interpretation and feature representation. By harnessing the power of LLMs, this study explores a novel and complementary tool for PDAC biomarker discovery and early detection, paving the way for further advancements in transcriptomic analysis.

## MATERIAL AND METHODS

In order to provide a clear overview of our methodological approach, we present Figure 1, which illustrates the key steps involved in our analysis. This figure outlines the process of utilizing a transcriptome profiling microarray (TPM) expression dataset obtained from the Gene Expression Omnibus (GEO), followed by the feature selection phase aimed at identifying genes predictive of pancreatic ductal adenocarcinoma (PDAC) samples^17^. Post feature selection, these values were used to generate peptide sequences that were used to train LLMs. This visual representation serves to summarize the workflow and key components of our study, facilitating a better understanding of our analytical strategy.

**Figure 1.**
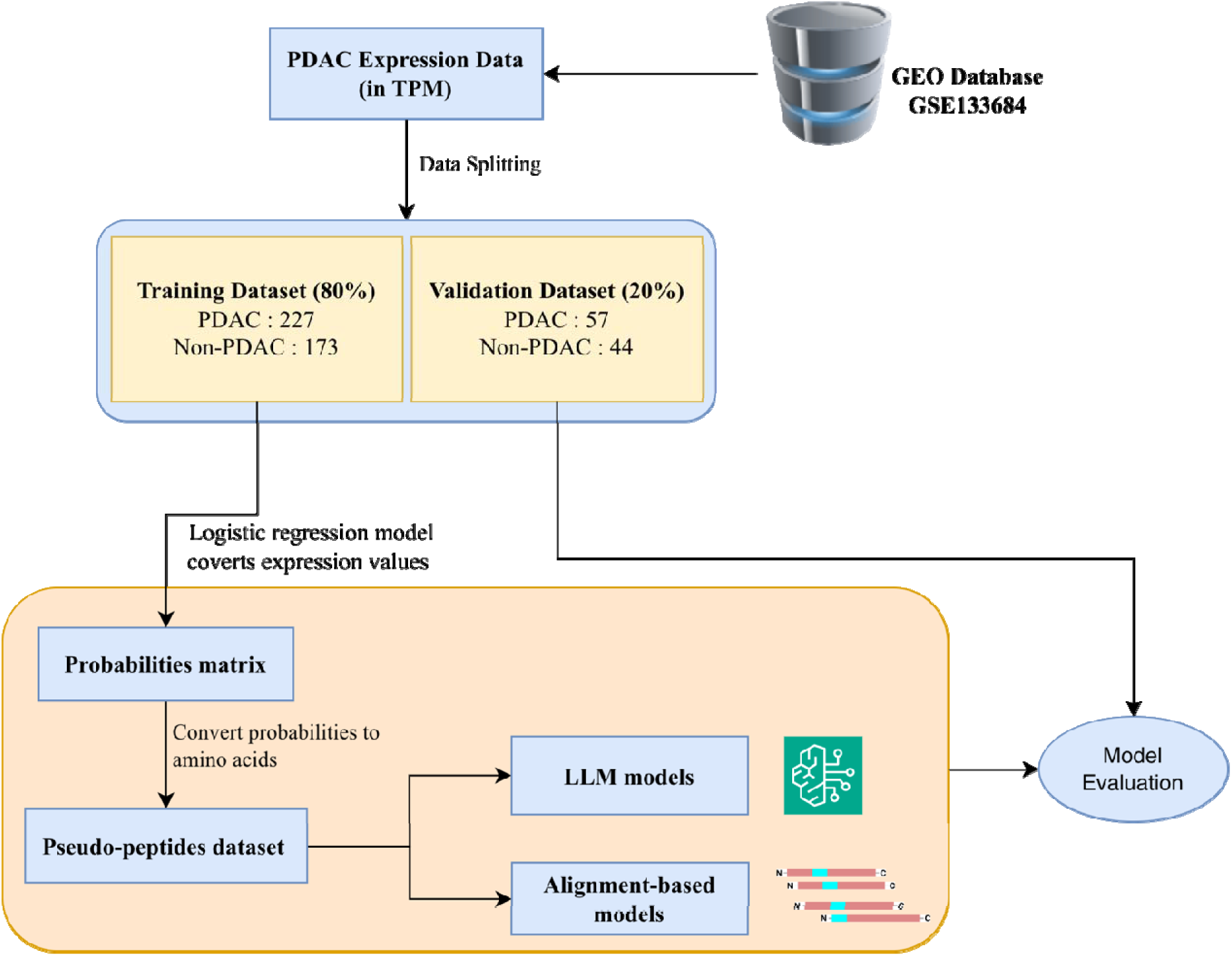
Overall methodology

### Dataset creation

Publicly available sequencing data was obtained from GEO (accession number: GSE133684) which was used in the study by Wang et al. Read counts were converted to TPM values by the depositors and the same was used for our study. The dataset consisted of 501 patient samples, which included - 284 PDAC patients, 100 Chronic Pancreatitis (CP) patients and 117 Healthy Controls (HC). The number of genes reported in this dataset was 54,148. For our analysis, labels were generated for each sample – with the positive label being PDAC samples (284 samples) and the negative label for CP and HC samples (217 samples). Subsequently the entire dataset was split into 2 parts - 80% in the training dataset (401 samples) and 20% in the validation dataset (100 samples).

### Expression to Probability

Following the preparation of the dataset, the TPM values were transformed into a probability matrix of the same dimensions (501 samples × 54,148 genes) to serve as features for subsequent analysis. This conversion was achieved by training a logistic regression (LR) model on each gene individually using the training dataset (401 samples). For each gene, the LR model was fitted to predict the binary outcome (PDAC vs. non-PDAC) based on its TPM values, and then applied to both the training and validation datasets to generate probability scores. This process was iteratively performed for all 54,148 genes, resulting in a probability matrix where each entry represents the likelihood of a sample being classified as PDAC based on the expression of a single gene. These probability values were then utilized as input features for the next stage of our analysis.

### Feature Selection

In order to reduce the high-dimensional feature space, originally comprising 54,148 features, a diverse set of eight feature selection methods was employed. All features were standardized using Z-score normalization prior to the selection process as the range for gene expression values for each gene has high variations. Each method was utilized to generate distinct feature subsets of nine specific sizes: 5, 8, 10, 15, 20, 25, 30, 40, and 50.

The first category included embedded regularization methods, which incorporate feature selection directly into the model training process. LASSO (L1 Regularization), implemented using LassoCV from sklearn, was applied. This linear model adds a penalty proportional to the absolute value (L1 norm) of the coefficients, promoting sparsity by forcing the coefficients of less important features to exactly zero. Complementing this, Elastic Net, implemented via ElasticNetCV, was also used. This method combines L1 and L2 penalties, making it particularly effective for selecting groups of correlated features where LASSO might arbitrarily select only one.

The second category consisted of tree-based importance methods, which rank features based on their contribution to the performance of ensemble models. Random Forest Importance was extracted from a RandomForestClassifier, utilizing the mean decrease in impurity (Gini importance) provided by each feature across all trees in the forest. Additionally, LGBM Importance was derived from an LGBMClassifier, a gradient-boosting framework that provided an alternative feature ranking based on criteria such as feature usage or gain.

Filter methods, which assess features based on their intrinsic statistical properties relative to the target variable, formed the third category. Information Gain was calculated using mutual_info_classif, measuring the mutual information between each feature and the target variable to quantify the predictive information one provides about the other. The ReliefF algorithm was also applied, implemented using the skrebate library. This instance-based method estimates feature quality by evaluating how well a feature’s value distinguishes between its nearest neighbors of the same and different classes.

The final category was wrapper methods, which use a specific machine learning model to "wrap" the selection process and iteratively evaluate feature subsets. Recursive Feature Elimination (RFE) with SVM was employed, using a Support Vector Machine (SVM) with a linear kernel as the base estimator. This process recursively fits the model, ranks features, discards the least important ones, and re-fits on the remaining set until the target number of features is reached. This same RFE process was repeated using a Random Forest classifier as the base estimator, leveraging its ensemble nature to guide the feature elimination.

All eight selection methods were implemented in Python using the sklearn, lightgbm, and skrebate libraries. The resulting feature sets for every method and size combination were systematically collected and saved to a JSON file for subsequent evaluation to identify the optimal feature set.

### Textual representation

A probability matrix was generated from patient gene expression values, where rows represent samples and columns correspond to genes. To generate sample-specific peptides, each probability value for all the genes selected was converted into an amino acid based on predefined probability ranges. Specifically, probabilities between 0 and 0.05 were mapped to alanine (A), 0.05 to 0.10 to cysteine (C), and so forth, following a systematic mapping scheme. A sample conversion of probabilities to peptides is shown graphically in Figure 2. For each sample, the amino acids derived from the probability values across all genes were concatenated to produce a unique peptide sequence for each sample. This process was repeated for all samples, resulting in a set of sample-specific peptide sequences reflective of their gene expression profiles.

**Figure 2.**
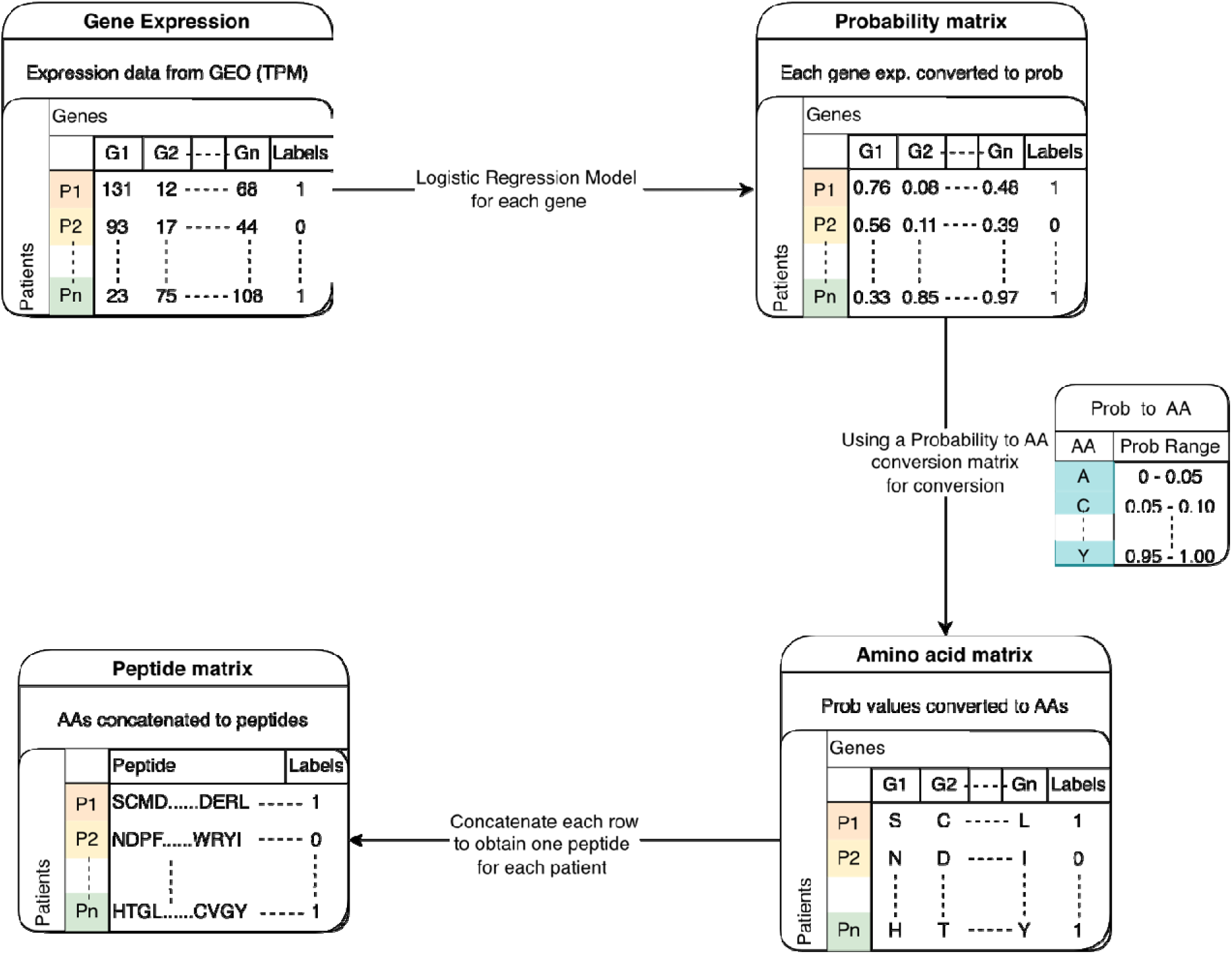
A graphical representation of the process for generating a representative peptide sequence for each patient gene expression data.

### Large Language Models

To classify PDAC patients based on the generated sample-specific peptide sequences, we employed several large language models (LLMs), including PeptideBERT, ProtBERT, t6_8M_UR50D, t12_35M_UR50D, and t33_650M_UR50D^18–20^. Each model was fine-tuned on the peptide sequences derived from our probability matrix, enabling the extraction of relevant features that capture the biological significance of the peptides. The training process involved using the training dataset (401 samples) to optimize the models for distinguishing PDAC patients from non-PDAC samples. Following training, we evaluated the performance of each model on the validation dataset (100 samples) using metrics such as accuracy, precision, recall, and F1- score. This comprehensive approach aimed to identify the most effective LLM for classifying PDAC patients, thereby enhancing our understanding of the molecular signatures associated with pancreatic cancer.

### Alignment-based methods

In this study, alignment-based methods were employed to classify pseudo-peptides of length-50 using PSI-BLAST and the MERCI motif search tool^21,22^. For the PSI-BLAST analysis, a database was created using 80% of the training dataset, and sequences from the validation dataset were queried against this database. The label of the best-matching sequence in the database was assigned to the query pseudo-peptide. Concurrently, the MERCI tool was used to identify motifs specific to both the positive and negative classes in the training data. These class-specific motifs were then applied to classify the validation sequences: if a positive-class motif was detected in a validation sequence, it was assigned a positive label, whereas the presence of a negative-class motif resulted in a negative label.

### Evaluation metrics

The binary classification performance of our fine-tuned model was evaluated using the following metrics: Sensitivity (SENS), Specificity (SPEC), Precision (PREC), Accuracy (ACC), Matthew’s Correlation Coefficient (MCC), F1-Score (F1) and Area Under the Receiver Operator Characteristic curve (AUC). The aforementioned metrics were calculated using the four different types of prediction outcomes: true positive (TP), false positive (FP), true negative (TN), and false negative (FN):

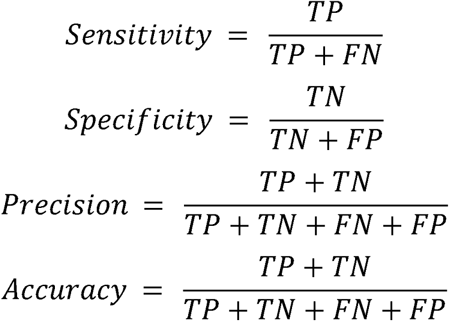

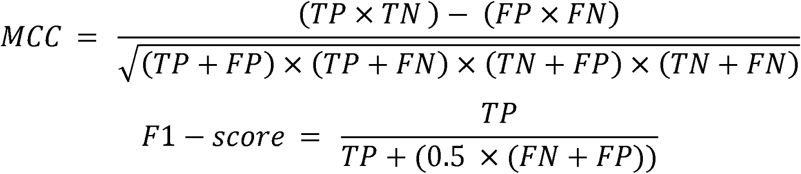

The evaluation of our binary classification model using various metrics provides critical insights into its performance. Sensitivity (SENS) measures the model’s ability to identify positive instances, while Specificity (SPEC) assesses its accuracy in recognizing negative instances. Precision (PREC) reflects the accuracy of positive predictions, and Accuracy (ACC) offers an overall measure of correctness, though it may be misleading in imbalanced datasets. Matthew’s Correlation Coefficient (MCC) provides a balanced view by considering all prediction outcomes, with values close to 1 indicating strong predictive capability. The F1-Score (F1) combines Precision and Sensitivity into a single metric, ideal for balancing the trade-off between false positives and negatives. Finally, the Area Under the Curve (AUC) evaluates the model’s ability to distinguish between classes across different thresholds, with higher values indicating better performance. Together, these metrics enable a comprehensive evaluation of the model, guiding necessary improvements and refinements.

### Ensemble methods

In this ensemble approach, the prediction probabilities from LLMs were combined with motif information from MERCI and sequence similarity data obtained through PSI-BLAST to provide a more accurate and biologically meaningful prediction of PDAC class peptide. Similar to how motifs from MERCI were incorporated, the query peptide was compared against the test data using PSI-BLAST, with varying e-values. If any motif or similarity hits were detected in the peptide, the probability score for the peptide was adjusted: for positive class peptides, the score was increased by 0.5, while for negative class peptides, the score was decreased by 0.5. This adjustment allowed the model to output a probability score, reflecting the likelihood of a peptide belonging to a specific class rather than a simple binary outcome.

## RESULTS

In this research, we aimed to develop a predictive model using a large language model (PeptideBERT) that is capable of identifying phenotypes of pancreatic ductal adenocarcinoma (PDAC) by analyzing gene expression data derived from blood-based exosomes. To achieve this objective, we utilized expression data obtained from a publicly accessible Gene Expression Omnibus (GEO) dataset. This dataset was generated using a non-invasive method for extracting RNA from blood-based exosomes, which were subsequently sequenced to gather the necessary genetic information.

### Selection of relevant genes

In this study, we aimed to identify discriminative genes that could serve as potential biomarkers for pancreatic ductal adenocarcinoma (PDAC). To achieve this, we applied multiple feature selection strategies to the gene expression dataset, with particular emphasis on the Light Gradient Boosting Machine (LGBM) model. The LGBM algorithm was utilized to rank genes according to their feature importance scores, thereby identifying those most strongly associated with PDAC classification. Based on this ranking, the top five genes—CLDN1, IL7R, ITIH2, KRT19, and MBNL1—were selected for further analysis due to their high discriminative potential and biological relevance.

Each of these genes has a well-supported mechanistic role in PDAC: CLDN1 promotes tumor growth, chemoresistance, and invasiveness through modulation of cell junctions and metabolic pathways^23^; IL7R is involved in regulating immune cell infiltration within the tumor microenvironment, which can influence patient prognosis^24^; ITIH2 acts as a metastasis suppressor by inhibiting cell motility and focal adhesion kinase signaling—its loss is linked to increased PDAC aggressiveness^25^; KRT19, a ductal marker, not only serves as a key biomarker for diagnosis but also facilitates immune evasion and is associated with poor prognosis^26^; and MBNL1, an RNA-binding protein, may restrict metastatic progression by governing alternative splicing and stabilizing tumor-suppressive transcripts^27^. The comparative analysis of their average expression levels between PDAC and non-PDAC samples is presented in Figure 3, demonstrating clear differential expression patterns. The complete ranked gene list derived from the LGBM analysis is provided in Supplementary Table S1 and the gene list for all the methods is provided in Supplementary Table S2.

**Figure 3.**
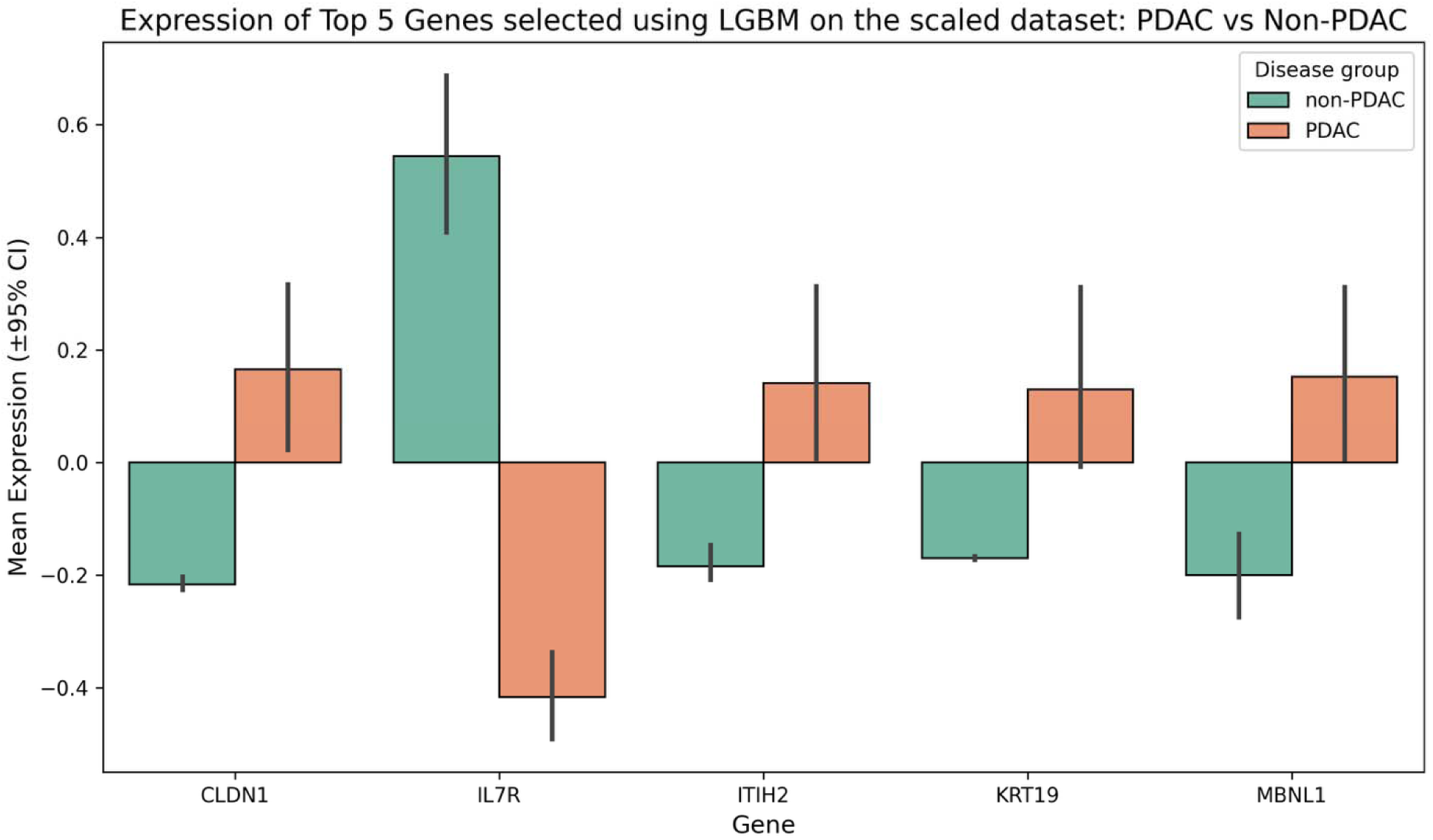
Bar plot highlighting the difference in the average expression of the genes selected by the LGBM algorithm between PDAC and non-PDAC samples; the dataset was scaled using Z-score normalization.

### Performance of LLMs

In this study, we employed a set of generated peptides to train several large language model (LLMs)—PeptideBERT, ProtBert, t6_8M_UR50D, t12_35M_UR50D, and t33_650M_UR50D - for the classification of pancreatic ductal adenocarcinoma (PDAC) patients. The models were trained on peptide sequences derived from PDAC patient samples to differentiate between diseased and healthy states, with performance evaluated using the area under the receiver operating characteristic curve (AUC). Among the ESM2-based models, t12_35M_UR50D achieved the highest AUC of 0.941, followed closely by t6_8M_UR50D (AUC = 0.936) and t33_650M_UR50D (AUC = 0.914), suggesting robust discriminatory power influenced by their pre-training on the UR50D dataset. However, the best overall performance was observed for PeptideBERT and ProtBert, where the models consistently outperform other approaches with an AUC of 0.942 and 0.962, respectively. The detailed comparison between the LLM models on the 50-length dataset is provided in Table 1. Table 2 shows the AUC values for all the LLM models corresponding to each length of peptide used in the study. It was also observed that PeptideBERT provides more accurate predictions for the smaller peptide lengths, while ProtBERT works better for longer peptide sequences. These results underscore the effectiveness of peptide-trained LLMs for PDAC classification and support further exploration of model optimization for clinical applications. The detailed performance of all the LLMs for different peptide lengths is provided in Supplementary Table S3.

**Table 1.** Performance of LLMs on the validation dataset for peptide length 50.

| Model | Sensitivity | Specificity | Precision | Accuracy | MCC | F1-score | AUC |
| --- | --- | --- | --- | --- | --- | --- | --- |
| Peptidebert | 0.929 | 0.613 | 0.757 | 0.792 | 0.584 | 0.834 | 0.942 |
| Protbert | **0.894** | **0.863** | **0.894** | **0.881** | **0.758** | **0.894** | **0.962** |
| t6_8M_UR50D | 0.877 | 0.818 | 0.862 | 0.851 | 0.697 | 0.869 | 0.936 |
| t12_35M_UR50D | 0.894 | 0.818 | 0.864 | 0.861 | 0.717 | 0.879 | 0.941 |
| t33_650M_UR50D | 0.842 | 0.818 | 0.857 | 0.8316 | 0.658 | 0.849 | 0.914 |

**Table 2.** AUC of LLMs on the validation dataset for peptide lengths - 5, 10, 15, 20, 25, 30, 40, 50.

| Model | k=5 | k=8 | k=10 | k=15 | k=20 | k=25 | k=30 | k=40 | k=50 |
| --- | --- | --- | --- | --- | --- | --- | --- | --- | --- |
| Peptidebert | **0.903** | **0.918** | 0.928 | 0.955 | **0.961** | **0.951** | 0.95 | 0.951 | 0.942 |
| Protbert | 0.825 | 0.824 | 0.918 | **0.957** | 0.957 | 0.905 | **0.957** | **0.961** | **0.962** |
| t6_8M_UR50D | 0.873 | 0.851 | 0.901 | 0.91 | 0.945 | 0.941 | 0.941 | 0.934 | 0.936 |
| t12_35M_UR50D | 0.870 | 0.865 | 0.932 | 0.913 | 0.935 | 0.934 | 0.944 | 0.939 | 0.941 |
| t33_650M_UR50D | 0.879 | 0.847 | **0.939** | 0.940 | 0.945 | 0.931 | 0.914 | 0.923 | 0.914 |

### Performance of Alignment based method

PSI-BLAST was used to perform a similarity search against a training dataset consisting of pseudo-peptides of length 50. The search was conducted with PSI-BLAST using an e-value range from 10^-7 to 10. Optimal performance was achieved at an e-value of 10^-3, correctly identifying 36 pseudo-peptides of the PDAC class and 9 pseudo-peptides from the non-PDAC class, with 9 incorrect hits. E-values lower than 10^-3 resulted in inadequate sequence coverage, while higher e-values led to a higher error rate. For peptide sequences of length 5, no hits were obtained while performing a BLAST query. The performance of this approach for peptide length 50 is shown in Table 3.

**Table 3.** Results for the BLAST-based alignment approach on the validation dataset.

| E-value | T_hits | C_hits | I_hits | P_hits | N_hits | CP_hits | CN_hits |
| --- | --- | --- | --- | --- | --- | --- | --- |
| 1.00E+03 | 101 | 80 | 21 | 57 | 44 | 51 | 29 |
| 1.00E+02 | 101 | 80 | 21 | 57 | 44 | 51 | 29 |
| 1.00E+01 | 101 | 80 | 21 | 57 | 44 | 51 | 29 |
| 1.00E+00 | 97 | 77 | 20 | 57 | 40 | 51 | 26 |
| 1.00E-01 | 89 | 70 | 19 | 52 | 37 | 46 | 24 |
| 1.00E-02 | 74 | 60 | 16 | 45 | 29 | 41 | 19 |
| 1.00E-03 | 54 | 45 | 9 | 37 | 17 | 36 | 9 |
| 1.00E-04 | 40 | 35 | 5 | 31 | 9 | 30 | 5 |
| 1.00E-05 | 35 | 31 | 4 | 30 | 5 | 29 | 2 |

Similarly, we used the MERCI tool for motif search, testing various fp and k values. fp and k values indicate the minimum hits of a motif in the positive class required to be considered as a motif and the number of motifs to be identified, respectively. Among them, fp 10 was chosen as the optimal parameter due to its high accuracy, even though it offered limited coverage. The result of the motif search is provided in Table 4.

**Table 4.** Results for the MERCI-based alignment approach on the validation dataset for peptides of length 50.

| **K** | **T_hits** | **C_hits** | **I_hits** | **P_hits** | **N_hits** | **CP_hits** | **CN_hits** |
| --- | --- | --- | --- | --- | --- | --- | --- |
| **10** | 28 | 27 | 1 | 27 | 5 | 26 | 2 |
| **100** | 30 | 29 | 1 | 29 | 1 | 28 | 1 |
| **1000** | 30 | 29 | 1 | 29 | 1 | 28 | 1 |

### Performance of Ensemble models

To further enhance the predictive performance of our models, we implemented ensemble methods that combined the predicted probabilities from the various large language models (LLMs) trained on peptide sequences with alignment-based methods. Specifically, we utilized both the MERCI algorithm and BLAST to integrate the predictions from PeptideBERT, ProtBert, t6_8M_UR50D, t12_35M_UR50D, and t33_650M_UR50D. By combining the predicted probabilities with the sequence similarity information provided by these methods, we aimed to refine our classification of PDAC patients and leverage the strengths of both the LLMs and the alignment-based approaches. The integration of predictions with MERCI resulted in a significant improvement in classification accuracy as compared to BLAST, demonstrating the effectiveness of combining predictive modeling with sequence alignment techniques. The performance of LLMs combined with BLAST and MERCI on the validation dataset is reported in Table 5. The detailed performance metric for the ensemble models for BLAST and MERCI is provided in Supplementary Tables S4 and S5, respectively.

**Table 5.** AUC of LLM + BLAST (BL) / MERCI (MR) ensemble models on the validation dataset for different peptide lengths (k)

| **Model** | **k=8** | **k=10** | **k=15** | **k=20** | **k=25** | **k=30** | **k=40** | **k=50** |
| --- | --- | --- | --- | --- | --- | --- | --- | --- |
| **Peptidebert** | 0.918 | 0.928 | 0.955 | **0.961** | **0.951** | 0.950 | 0.951 | 0.942 |
| **Peptidebert + BL** | 0.918 | 0.934 | 0.955 | 0.956 | 0.944 | 0.945 | 0.952 | 0.932 |
| **Peptidebert + MR** | **0.925** | 0.932 | 0.949 | 0.954 | 0.950 | 0.946 | 0.952 | 0.942 |
| **Protbert** | 0.824 | 0.918 | **0.957** | 0.957 | 0.905 | **0.957** | **0.961** | **0.962** |
| **Protbert + BL** | 0.823 | 0.922 | 0.956 | 0.955 | 0.904 | 0.956 | 0.963 | 0.951 |
| **Protbert + MR** | 0.874 | 0.932 | 0.950 | 0.957 | 0.776 | 0.954 | 0.966 | 0.960 |
| **t6_8M_UR50D** | 0.851 | 0.901 | 0.910 | 0.945 | 0.941 | 0.941 | 0.934 | 0.936 |
| **t6_8M_UR50D+ BL** | 0.850 | 0.905 | 0.912 | 0.937 | 0.940 | 0.942 | 0.938 | 0.927 |
| **t6_8M_UR50D+ MR** | 0.870 | 0.908 | 0.914 | 0.944 | 0.944 | 0.939 | 0.936 | 0.935 |
| **t12_35M_UR50D** | 0.865 | 0.932 | 0.913 | 0.935 | 0.934 | 0.944 | 0.939 | 0.941 |
| **t12_35M_UR50D+ BL** | 0.864 | 0.934 | 0.913 | 0.930 | 0.924 | 0.945 | 0.941 | 0.929 |
| **t12_35M_UR50D+ MR** | 0.873 | 0.926 | 0.908 | 0.934 | 0.935 | 0.941 | 0.939 | 0.941 |
| **t33_650M_UR50D** | 0.847 | **0.939** | 0.940 | 0.945 | 0.931 | 0.914 | 0.923 | 0.914 |
| **t33_650M_UR50D+ BL** | 0.846 | 0.938 | 0.940 | 0.944 | 0.928 | 0.913 | 0.931 | 0.915 |
| **t33_650M_UR50D+ MR** | 0.863 | 0.936 | 0.935 | 0.947 | 0.932 | 0.912 | 0.928 | 0.919 |

### Comparison with the existing methods

We compared the performance of our ensemble models with two existing methods that have previously attempted to classify pancreatic ductal adenocarcinoma (PDAC) based on blood exosome derived transcriptomics. Wang et al., which utilizes a SVM model to classify PDAC patients based on the expression of an eight gene biomarker signature, managed to achieve an AUC of 0.936 on their own external validation dataset. A method by Wang et al., identified a four lncRNA signature that could classify PDAC patients with an AUC of 0.848. Our ensemble models, which integrated predictions from multiple large language models with alignment-based methods, consistently outperformed both the tools. The detailed comparison of the existing methods with our approach is reported in Table 6. The performance of the existing tools represents the values that were reported in their respective studies. Notably, the AUC values for our models were significantly higher, indicating superior discriminative ability in classifying PDAC patients. This comparison highlights the advantages of our approach, which leverages the strengths of both LLMs and alignment techniques, ultimately providing a more robust and accurate tool for clinical applications in PDAC detection and management.

**Table 6.** Comparison of performance reported by existing methods with our method for peptide lengths 5, 8 and 10.

|  | Number of gene in biomarker panel | Sensitivity | Specificity | Precision | Accuracy | MCC | F1-score | AUC |
| --- | --- | --- | --- | --- | --- | --- | --- | --- |
| Wang et al. 2020 | 8 | 0.937 | 0.916 | - | 0.927 | - | - | 0.936 |
| Wang et al. 2024 | 4 | 0.72 | 0.89 | - | - | - | - | 0.848 |
| Peptidebert | 5 | 0.825 | 0.841 | 0.87 | 0.832 | 0.662 | 0.847 | 0.903 |
| Peptidebert + MR | 8 | 0.912 | 0.614 | 0.754 | 0.782 | 0.561 | 0.825 | 0.925 |
| t33_650M_UR50D | 10 | 0.895 | 0.886 | 0.911 | 0.891 | 0.779 | 0.903 | 0.939 |

## DISCUSSION

This study marks a significant progress in the application of artificial intelligence (AI) and machine learning (ML) for pancreatic ductal adenocarcinoma (PDAC) diagnosis by introducing large language models (LLMs) as a novel methodology for leveraging transcriptomic data. By transforming gene expression profiles into pseudo-peptides and training a PeptideBERT model, we achieved a robust classification performance with an AUC of 0.939. This outcome underscores the transformative potential of LLMs in bioinformatics, particularly in translating complex transcriptomic data into interpretable and actionable insights for disease prediction.

The unique ability of LLMs to process and model sequential data positions them as a powerful tool for analyzing biological sequences. In this study, the transformation of gene expression data into a textual format proved pivotal, enabling the PeptideBERT model to identify meaningful patterns within the data. This approach not only demonstrated comparable or superior performance to traditional methodologies but also highlighted the scalability of LLMs in analyzing high-dimensional biological datasets.

Moreover, this study broadens the horizon of LLM applications in precision oncology, providing a robust framework for integrating diverse types of omics data for diagnostic purposes. The versatility of LLMs in handling complex sequential data, as demonstrated here, represents a paradigm shift in how transcriptomic data can be used for early and accurate disease detection. The results of this study establish a strong foundation for future exploration of LLMs in precision diagnostics, including their potential for analyzing data beyond transcriptomics, such as proteomics or metabolomics.

However, there are still certain limitations of the study that can be addressed in the future. In the process of generating the peptide sequences, the position of the amino acids is determined by the probabilities and the peptides generated from them do not represent real world peptides. This could mislead pattern recognition in LLMs which are trained on real peptides. The length of the generated peptides in our study is limited to 50, whereas it may be possible that longer peptides may report better accuracies in identifying PDAC samples.

## CONCLUSION

While this study highlights the potential of large language models (LLMs) in improving precision diagnostics for pancreatic ductal adenocarcinoma (PDAC), certain challenges such as computational complexity and resource demands must be addressed for clinical implementation. Optimizing LLMs for efficiency and scalability is crucial to their adoption in routine diagnostics.

Additionally, as LLMs evolve, their integration with multi-omics data and predictive modelling for treatment response offers significant promise. These advancements could transform PDAC diagnosis and management, paving the way for more accurate, comprehensive, and personalized healthcare solutions.

## ACKNOWLEDGEMENTS

Authors are thankful to the Department of Science and Technology (DST-INSPIRE) and Indraprastha Institute of Information Technology, New Delhi, for fellowships and financial support. Authors are also thankful to Department of Computational Biology, IIITD New Delhi for infrastructure and facilities.

## AUTHORS’ CONTRIBUTIONS

SC collected and processed the datasets. SC and NKM implemented the algorithms and developed the prediction models. SC, NKM and GPSR analyzed the results. SC, NKM and GPSR penned the manuscript. GPSR conceived and coordinated the project. All authors have read and approved the final manuscript.

